# The Venification of the Artery: Insights from Single-Cell Transcriptomics of a Porcine Arteriovenous Fistula Model

**DOI:** 10.64898/2026.09.23.753931

**Authors:** Matthew S. Sussman, Marwan Tabbara, Xochilt Labissiere, Juan S. Lopez-McCormick, Santiago J. Guzman, Christina Kosanovic, Roberto I. Vazquez-Padron, Laisel Martinez

**Author notes:** **Correspondence:** Laisel Martinez, PharmD, MS, Division of Vascular and Endovascular Surgery, University of Miami Miller School of Medicine, 1600 NW 10^th^ Avenue, RMSB 1042, Miami, FL 33136, **E-mail:**. **What this paper adds:** The study provides a holistic view of the hemodynamic adaptation of autologous arteriovenous fistulas (AVF) by integrating histologic and molecular analyses of both the arterial and venous limbs. Using single-cell transcriptomics, we demonstrate at cellular resolution that vascular adaptation in AVFs extends beyond the well-studied concept of venous arterialization and includes an even more prominent process of arterial venification. By defining both the shared and vessel-specific mechanisms driving vascular remodeling, these findings may inform the development of vascular-wide and vessel-selective therapeutic strategies to improve the maturation and patency of vascular accesses.

## Abstract

**Objective:** The maturation of an autologous arteriovenous fistula (AVF) depends on the coordinated hemodynamic adaptation of the inflow artery and the outflow vein. While venous remodeling has been extensively studied, the mechanisms of arterial adaptation are poorly understood. We addressed this knowledge gap with the first temporal single-cell atlas of the porcine AVF model.

**Methods:** Six Yorkshire pigs underwent bilateral femoro-femoral AVF creation for tissue harvest at 2 and 21 days postoperatively or sham operation to collect native vessels after 2 days. Arteries and veins were analyzed by droplet-based single-cell RNA sequencing, histology, and immunohistochemistry.

**Results:** All fistulas were patent, with flows reaching 1185.0 ± 170.6 mL/min despite mild to moderate neointima formation in the veins. Arteries and veins demonstrated >60% endothelial denudation and increased neovascularization. Arterial remodeling was characterized by a progressive increase in venous-like transcriptional features. Arteries presented a venous-like endothelial cell (EC) composition at 21 days postop and a temporal increase in venous-associated phenotypes of myofibroblasts and fibroblasts. In both vessels, the latter populations were the main bearers of mechanosensitive signatures at 2 days. Despite the temporal convergence in phenotypes, arteries and veins retained vessel-selective characteristics. Arterial ECs expressed high levels of nitric oxide synthase, whereas venous ECs were largely inflammatory. Arterial myofibroblasts and fibroblasts had higher expression of Notch signaling transducers (*HEY2, HEYL*) and cellular communication network factors (*CCN3, CCN5*), while Wnt signaling regulators (*DKK2, SFRP4*) and complement genes (*C3, C7, CFD*) were upregulated in veins. Anti-inflammatory macrophages increased in both vessels during remodeling, likely contributing to the maturation of the fistulas.

**Conclusion:** These analyses challenge the traditional paradigm of exclusive vein arterialization and support a complementary process of arterial venification after anastomosis. Recognition of this underappreciated phenomenon may reshape our understanding of AVF biology and guide the discovery of both vascular-wide and vessel-selective anti-stenotic therapies.

## INTRODUCTION

The multicenter Hemodialysis Fistula Maturation (HFM) Study reported 52% of unassisted maturation failure, with 37% of arteriovenous fistulas (AVF) requiring interventions to facilitate maturation, and 26% that ultimately failed overall maturation.^1^ Prevention of these early failures has been a priority for vascular surgeons, nephrologists, and basic scientists. Unfortunately, over 30 years of basic and translational research have yielded minimal improvements for this longstanding surgical problem. The failure of multiple clinical trials to improve AVF maturation^2–5^ has exposed fundamental limitations in our understanding of AVF biology, underscoring the need for a better assessment of biological models that explain the failure of adaptive remodeling in both the venous and arterial limbs of the AVF.^6, 7^

Historically, research on AVF maturation has focused almost exclusively on the vein, largely overlooking the equally important contribution of adaptive remodeling in the feeding artery. This emphasis is driven, in part, by the high incidence of venous stenoses, as well as the technical challenges associated with obtaining arterial biopsies from patients. Nonetheless, prospective observational studies have underscored the critical role of arterial function and adaptation in successful AVF maturation.^8–10^ Preoperative arterial diameter has been recognized as an important predictor of AVF remodeling.^11^ A *post hoc* HFM analysis also demonstrated a strong association of wall shear stress in the postoperative artery with unassisted maturation of the AVF.^9^

In this work, we generated the first high-resolution cellular atlas of the porcine femoro-femoral AVF, providing a unique view of the evolving cellular landscape in both the inflow artery and the outflow vein after anastomosis. Using this clinically relevant model, we define the molecular basis of venous arterialization but also present evidence in support of “arterial venification”, a rarely reported process in AVFs. Our analyses reveal how supraphysiologic hemodynamic forces override the preoperative differences between arteries and veins, inducing convergent remodeling mechanisms in both types of vessels that drive AVF maturation.

## MATERIALS AND METHODS

### Study Design

Six Yorkshire pigs, ∼50 kg in weight, were allocated to 1) bilateral femoro-femoral AVF creation for tissue harvest at 2 days postop, 2) bilateral AVFs for harvest at 21 days, or 3) bilateral sham operation to collect native vessels 2 days after opening and closing the skin and fascia (**Figure 1A**). Each experimental arm included one male and one female. Therefore, a total of four arteries and four veins were collected per group for histology and single-cell RNA sequencing (scRNA-seq) (**Supplementary Materials and Methods**).

**Figure 1.**
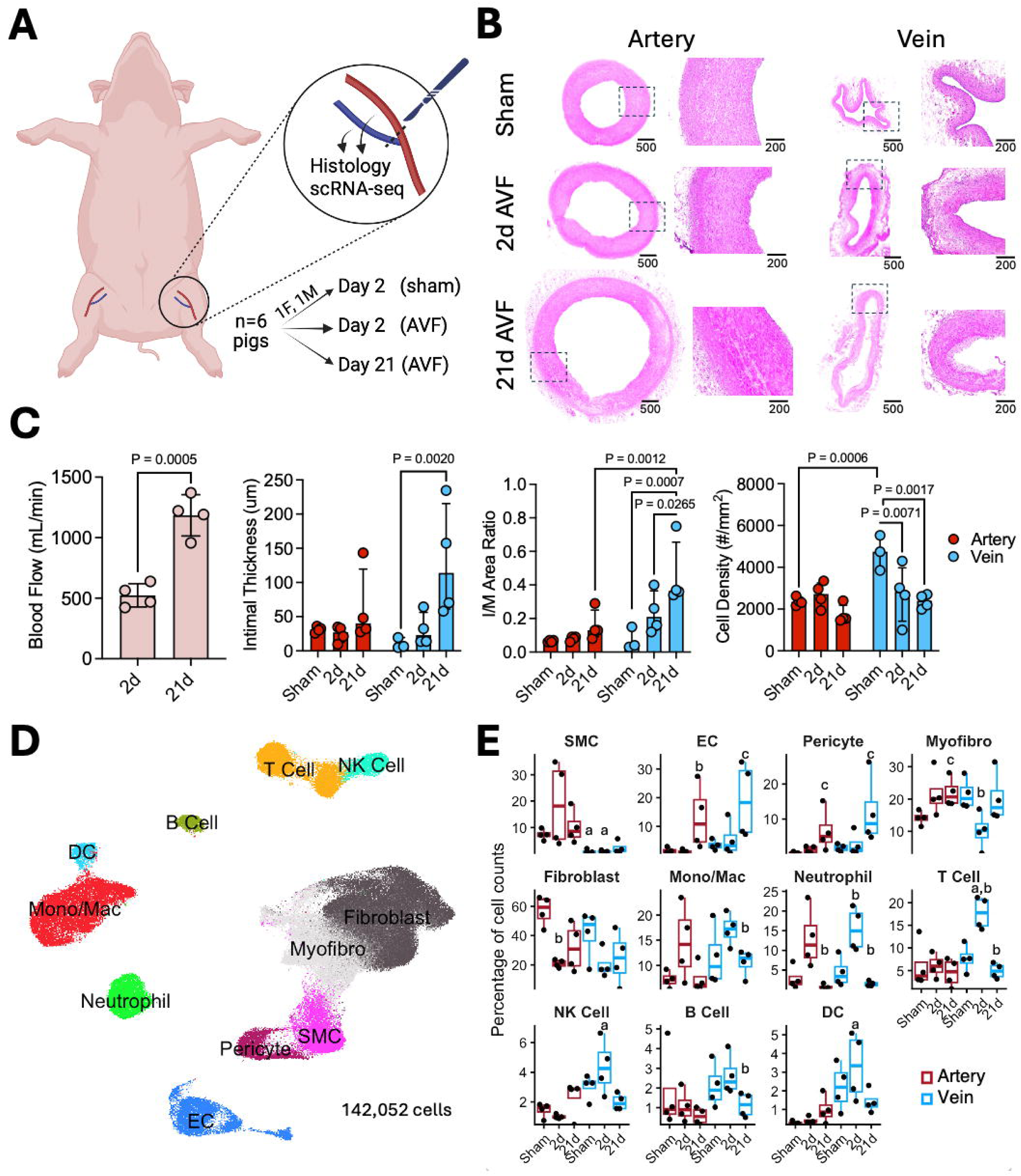
Arterial and venous remodeling in the pig femoro-femoral AVF model. **A)** Study design. **B)** Representative histology of arteries and veins per time point of tissue collection. Dashed boxes are magnified to the right. **C)** Functional assessment and histomorphometry of AVF remodeling. Error bars indicate the mean and standard deviation or the median and interquartile range as appropriate. Only significant p-values are shown. **D)** Integrated uniform manifold approximation and projection (UMAP) of 142,052 single cells from 12 femoral arteries and 12 femoral veins. **E)** Relative proportions of cell populations per sample. a, P<0.05 compared with the artery; b, P<0.05 compared with the previous time point within the same vessel type; c, P<0.05 compared with the sham group within the same vessel type.

### Surgical Procedures

All procedures were performed by board-certified vascular surgeons as described in the **Supplementary Materials and Methods**. Briefly, femoral vein to femoral artery AVFs were created using an ∼5 mm end-to-side anastomosis with 7-0 polypropylene suture. Animal experiments were approved by the University of Miami Institutional Animal Care and Use Committee.

### Single-Cell RNA Sequencing

Sequencing was performed at the University of Miami John P. Hussman Institute for Human Genomics (**Supplementary Materials and Methods**) and analyzed using published bioinformatic pipelines.^12–15^ Sequencing data were deposited in the Gene Expression Omnibus repository with accession number GSEXXXXXX.

### Histology and Immunostaining

Tissue sections were stained with hematoxylin and eosin for gross morphometric analysis. The protein expressions of CD31, CD11b, endothelial nitric oxide (NO) synthase (eNOS), and Ki-67 were assessed by immunohistochemistry (IHC) or immunofluorescence (**Supplementary Materials and Methods**).

### Statistical Analyses

Groups were compared using parametric or non-parametric tests based on data distribution assessed with the Shapiro-Wilk test (**Supplementary Materials and Methods**).

## RESULTS

### Physiological maturation of the swine AVF at the histologic and cellular levels

We analyzed the morphometry and cellular composition of arteries and veins in femoro-femoral AVFs using histology and scRNA-seq (**Figure 1A-B**). All AVFs were patent at the time of harvest, with mean blood flow increasing from 524 mL/min at 2 days to 1185 mL/min at day 21 postop (**Figure 1C**). The average wall thickness remained constant in arteries over time but increased progressively in veins (**Supplementary Figure S1**). Veins developed mild to moderate intimal hyperplasia (IH) at 21 days that was significantly higher than in arteries at the same time point. The cellular density decreased in veins after AVF creation reflecting proportionally higher extracellular matrix (ECM) deposition than cell accumulation (**Figure 1C**).

We generated an integrated map of 142,052 cells from the 12 arteries and 12 veins by scRNA-seq, with cells clusters manually annotated based on canonical markers^16, 17^ (**Figure 1D, Supplementary Figure S2**). The map included comparable numbers of cells from experimental groups and individual vessels (**Supplementary Figure S3**). Smooth muscle cells (SMC) were more abundant in arteries than in veins before and after anastomosis (**Figure 1E**). Arteries exhibited a progressive expansion of myofibroblasts that reached statistical significance at 21 days, likely at the expense of resident fibroblasts, whose abundance decreased 2 days after surgery. In contrast, veins showed an early reduction in myofibroblasts at 2 days, followed by a partial recovery by day 21. Both vessels had similar proportions of fibroblasts by day 21. Endothelial cells (EC) and pericytes increased in abundance at 21 days in both arteries and veins, suggesting a postoperative expansion of the intramural microvasculature (**Figure 1E**).

The proportions of immune cells after surgery uncovered unique characteristics of early immune cell mobilization in the vein. Arteries presented increasing trends in neutrophils and monocyte/macrophages 2 days after anastomosis. At this time in veins, there was a significant increment in neutrophils, an increasing trend in mono/macs, and significant increases of T cells, NK cells, and dendritic cells that were not observed in the arterial segment (**Figure 1E**).

### Expansion of the vasa vasorum and sustained luminal de-endothelialization

We analyzed the changes in EC phenotypes to better understand the arterial and venous responses to increased shear stress. Sub-clustering of 8,999 ECs identified seven distinct endothelial subtypes. These included the previously described venous, capillary, arteriolar, and lymphatic EC subpopulations (**Figure 2A-C**);^18^ two ECM-producing phenotypes termed elastogenic and pro-fibrotic, based on their differential expression of elastin (*ELN*) and collagen-maturation genes (e.g., *PCOLCE*); and a hemostatic EC population enriched in coagulation-related regulators (e.g., *PROCR*) that was abundant in the arteries early after anastomosis (**Supplementary Figure S4**).

**Figure 2.**
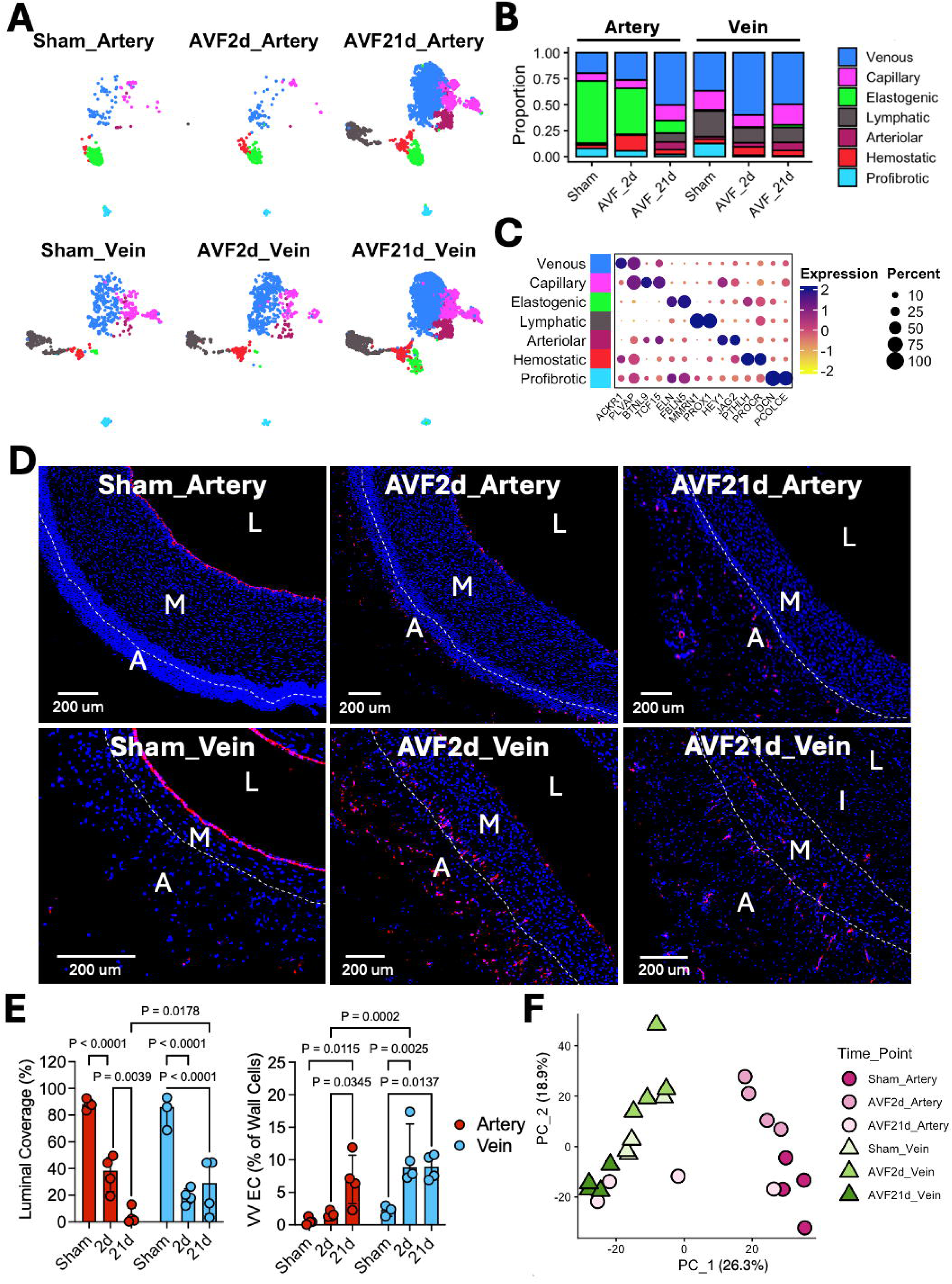
Convergence of arterial and venous endothelial cells after AVF creation. **A-B)** Focused UMAPs and mean cell proportions of endothelial cell (EC) subpopulations relative to the total number of ECs by experimental group. **C)** Dot plot representation of expression markers for EC subpopulations. Dark purple dots indicate the highest expression levels, while the size of the dot represents the percentage of cells within each subcluster expressing the gene. **D)** Representative immunofluorescence staining of the pan-EC marker CD31 per experimental group. Dash lines separate the vascular layers. L: lumen, I: intima, M: media, and A: adventitia. **E)** Quantification of CD31+ luminal coverage as percentage of luminal perimeter and of CD31+ vasa vasorum (VV) ECs as percentage of total DAPI+ cells. Error bars indicate the median and interquartile range. **F)** Principal component analysis of EC transcriptional variance at the sample level.

The compositional analysis of EC phenotypes revealed striking preoperative differences between arteries and veins that progressively diminished after surgery as the artery acquired venous-like characteristics. At baseline and 2 days postop, elastogenic ECs predominated in arteries (∼60%), but their abundance declined over time, consistent with significant luminal de-endothelialization beginning 2 days after surgery (**Figure 2B, D-E**). Sham and 2-day arteries also contained smaller proportions of venous, capillary, and arteriolar ECs, corresponding to sparse intramural microvessels (**Figure 2B**). By day 21, the arterial endothelium transitioned toward a venous-like EC composition, driven by postoperative expansion of the vasa vasorum (**Figure 2A–B, Supplementary Figure S4**).

In veins, the venous phenotype predominated before surgery (∼40%), while the capillary, lymphatic, and pro-fibrotic subtypes contributed 10-50% depending on the individual sample (**Supplementary Figure S4**). Veins also experienced significant luminal de-endothelialization 2 days after anastomosis, but maintained their EC composition due to the extensive number of venules in the vasa vasorum^17^ and a proportional increase in intramural vascularization (**Figure 2B, D-E**). The decrease in EC transcriptional variance between arteries and veins at 21 days was further illustrated by principal component analysis (PCA) (**Figure 2F**).

We next examined the genes that remained differentially expressed between arteries and veins in 21-day AVFs, and which may reflect vessel-specific microenvironments and responses to hemodynamic stimuli (**Figure 3A**). Genes upregulated in arterial subpopulations at this time point included positive (*BGN, MFAP5*)^19, 20^ and negative regulators (*MGP, IGFBP5*)^21, 22^ of angiogenesis in line with a controlled neovascularization, genes involved in elastogenesis (*ELN, LTBP2*),^23^ and eNOS (encoded by *NOS3*). Venous subpopulations, on the other hand, showed upregulation of potent angiogenic factors (*HGF*)^24^ and various inflammatory mediators (e.g., *IL6, IL33, CCL2, CXCL2*). These differences in expression were not unique to the 21-day time point. The major elastogenic population and minor hemostatic ECs of sham arteries, as well as remnants of these populations at 2 days, had the highest levels of *NOS3*, indicating a prominent role in shear stress sensing prior to denudation (**Figure 3B-C**). The hemostatic ECs of sham and 2-day veins also expressed *NOS3*, but at lower levels than arteries (**Figure 3B-C**). Similarly, high levels of inflammatory markers (e.g., *IL6* and *ICAM1*) were already present in venous subpopulations before surgery (**Figure 3B**).

**Figure 3.**
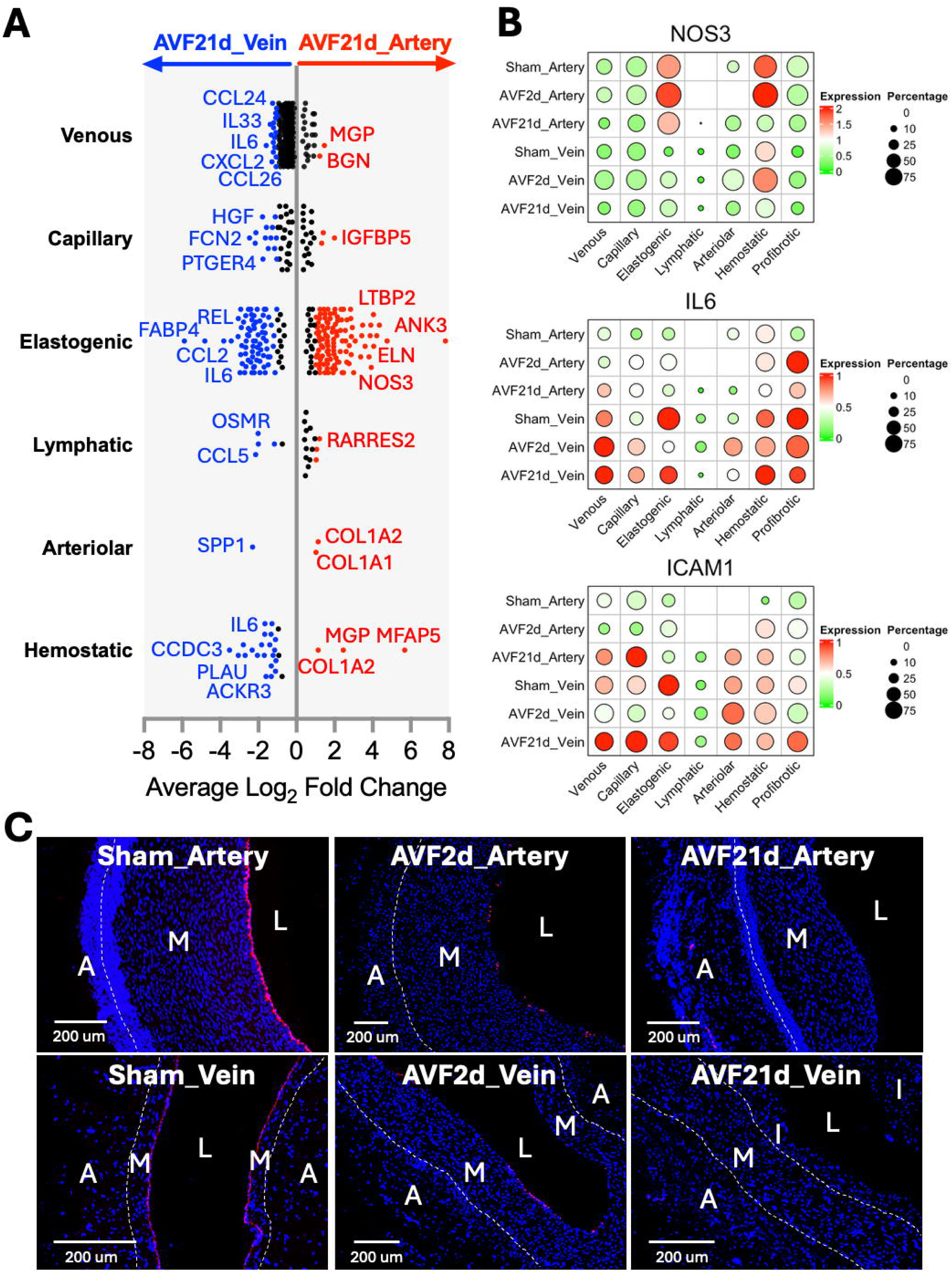
Unique microenvironments of arterial and venous endothelial cells. **A)** Differentially expressed genes (DEG) between arteries and veins from 21-day AVFs per endothelial cell (EC) subtype. Genes with FDR<0.01 and absolute log_2_(fold change [FC])>0.5 are plotted in the graph, but only those with log_2_FC>1 (up in arteries) or <-1 (up in veins) are colored in red and blue, respectively. **B)** Dot plot representation of selected DEGs per EC phenotype and experimental group. Bright red dots indicate the highest expression levels, while the size of the dot represents the percentage of cells expressing the gene. **C)** Representative immunofluorescence staining of endothelial nitric oxide synthase (eNOS) per experimental group. Dash lines separate the vascular layers. L: lumen, I: intima, M: media, and A: adventitia.

### Biomechanical sensing by fibroblastic cells and vessel-selective regulatory mechanisms

We then analyzed the mural cells (9,540 SMCs; 5,187 pericytes), 49,836 fibroblasts; and 26,654 myofibroblasts to identify signatures of mechanical stretching and wall remodeling. In addition to the differences in SMC proportions between vessel types and the increase in pericytes over time (**Figure 1E**), we found distinct subpopulations of myofibroblasts and fibroblasts that defined the early and late phases of remodeling (**Figure 4A-C**). Within the myofibroblast cluster, proliferative myofibroblasts were defined by cell division markers (*CCND1, NPM3*), pro-fibrotic myofibroblasts by high expression of periostin (*POSTN*), and inflammatory myofibroblasts by the expression of chemokines (e.g., *CXCL8*). Similarly, elastogenic fibroblasts were characterized by production of elastin and interacting proteins (*ELN, LTBP2*), secretory fibroblasts by complement factors (*CFD, C3*), and redox fibroblasts by the expression of anti-oxidative enzymes (*SOD2, MT1X*) (**Figure 4A-B**).

**Figure 4.**
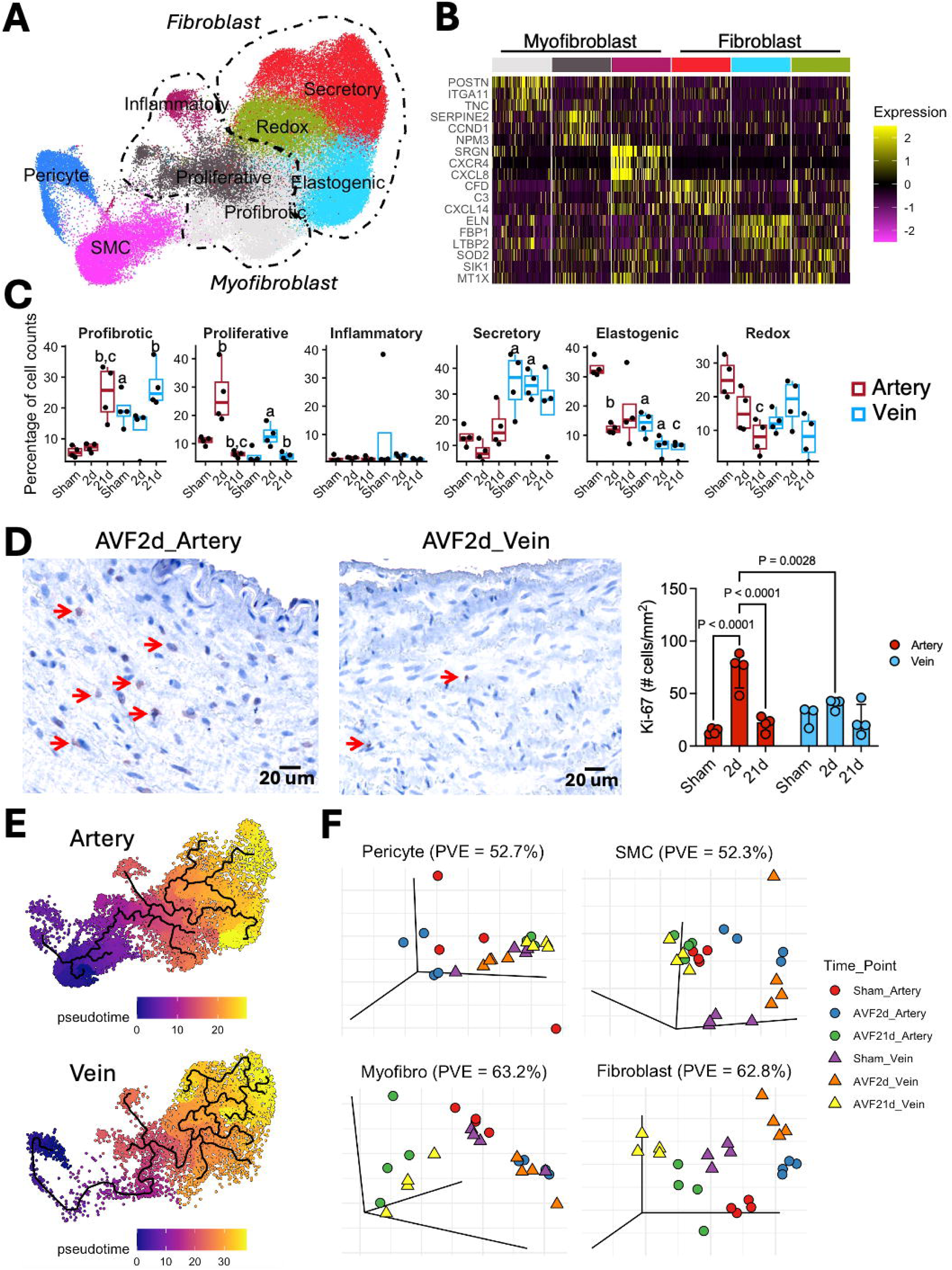
Convergent differentiation of fibroblastic cells in arteries and veins. **A)** Integrated UMAP of mural cells, myofibroblasts, and fibroblasts from 12 femoral arteries and 12 femoral veins, color-coded by cell phenotype. **B)** Heatmap representation of expression markers for subpopulations of myofibroblasts and fibroblasts, with bright yellow indicating the highest expression levels. **C)** Distribution of myofibroblast and fibroblast subtypes per sample relative to the total number of fibroblastic cells. a, P<0.05 compared with the artery; b, P<0.05 compared with the previous time point within the same vessel type; c, P<0.05 compared with the sham group within the same vessel type. **D)** Representative immunohistochemistry and quantification of Ki-67 staining in experimental groups. Arrows indicate positive cells. **E)** Pseudotime trajectory analysis of mural cells, myofibroblasts, and fibroblasts from 2-day arteries and veins. The trajectories start arbitrarily in mural cells. **F)** Principal component analyses of transcriptional variance in mural cells and fibroblastic cells at the sample level.

At baseline, femoral arteries contained a higher proportion of elastogenic fibroblasts than veins, whereas pro-fibrotic myofibroblasts and secretory fibroblasts were more abundant in the venous wall (**Figure 4C**). Early after anastomosis, myofibroblast proliferation was significantly greater in the arterial limb and correlated with the higher number of Ki-67+ cells by IHC (**Figure 4C–D**). Pseudotime trajectory analysis of the 2-day time point predicted that proliferating myofibroblasts in arteries arise directly from resident fibroblastic cells and SMCs, whereas the smaller proliferative population in veins is most closely related to existing myofibroblasts (**Figure 4E**). The late phase of remodeling coincided with a venous-like shift in the fibroblast composition of the arterial limb. Arteries exhibited a marked reduction in elastogenic fibroblasts, a gradual decline in redox fibroblasts, and an expansion of pro-fibrotic myofibroblasts that reached similar proportions as in veins by day 21 (**Figure 4C**). In contrast, secretory fibroblasts remained consistently more abundant in veins throughout the study, representing the only fibroblast subtype that did not converge between the two vessels. In PCAs, both mural cells and fibroblasts demonstrated a decrease in transcriptional variance between arteries and veins at 21 days (**Figure 4F**).

Given the parallel differentiation of arterial and venous cells, we paid attention to the transcriptional activation of mechanosensitive programs in each vascular bed. The propagation of mechanical clues to the cell nucleus relies on the family of Rho GTPases and kinases that modulate actin polymerization (e.g., RHOA, ROCK1, ROCK2), the anchoring of the cytoskeleton to the nuclear envelope (via emerin [EMD]), and activation of transcription factors such as MRTFs, YAP, and TAZ.^25–27^ In both arteries and veins, myofibroblasts and fibroblasts demonstrated robust activation of this machinery 2 days after anastomosis, although expression was higher in the arterial limb (**Figure 5A**). Similarly, in both vessels, mechanosensitive signatures were followed by the associated upregulation of ECM genes (e.g., *COL1A1, COL8A1*), cytoskeletal proteins (e.g., *ACTA2*), and mediators of cell migration and tissue remodeling (e.g., *FAP, POSTN, and CCN2*) by day 21 (**Figure 5A, Supplementary Figure S5**). In pericytes and SMCs, this temporal pattern was not as clear, possibly due to an earlier or continued upregulation of mechanosensitive genes (**Supplementary Figure S5**).

**Figure 5.**
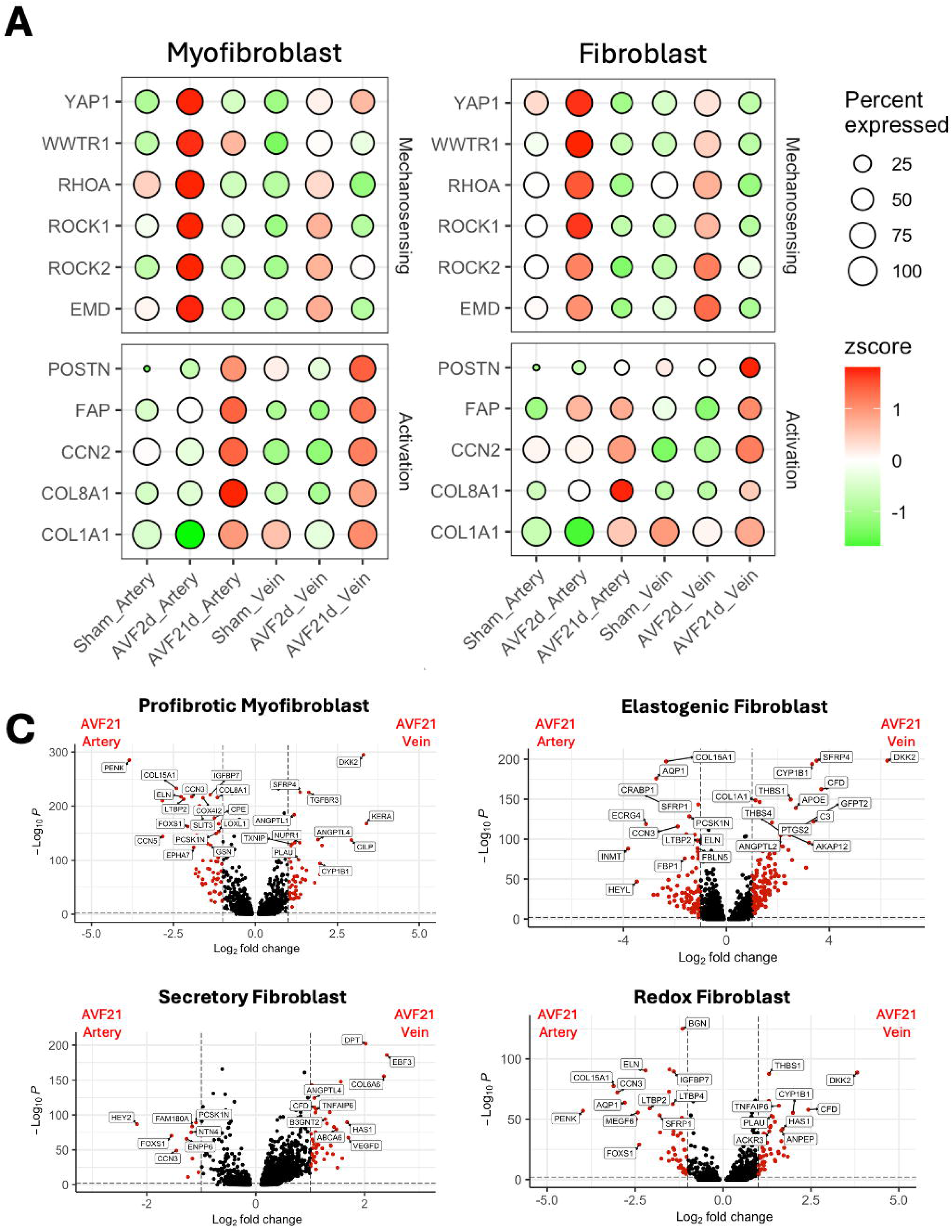
Mechanosensitive activation of fibroblastic cells from arteries and veins. **A)** Temporal upregulation of mechanotransduction genes and markers of fibroblastic cell activation in arteries and veins. The highest expression of mechanosensitive genes occurs in both vessels at 2 days postop, while markers of myofibroblast and fibroblast activation show upregulation at 21 days. Bright red dots indicate the highest expression levels, while the size of the dot represents the percentage of cells expressing the gene per experimental group. **B)** Volcano plots of differentially expressed genes (DEG; FDR<0.01 and absolute log_2_(fold change [FC])>1) between arteries and veins in the main fibroblastic cell populations at 21 days postop. Red dots indicate DEGs upregulated in arteries (log2FC<-1) or veins (log2FC>1).

Interestingly, several key regulators of fibroblast activation and myofibroblast differentiation remained differentially expressed between the two vessels independent of the time point (**Figure 5B, Supplementary Figure S6**). Upregulated genes in arteries included *AQP1*, a promoter of myofibroblast activation;^28^ *CCN3* and *CCN5*, both inhibitors of fibrosis,^29, 30^ as well as the Notch signaling transducers *HEY2* and *HEYL*, which determine the arterial cell fate.^31^ Compared with arteries, venous myofibroblasts and fibroblasts had higher expression of Wnt signaling regulators (*DKK2, SFRP4*) and complement genes (*C3, C7, CFD*).

### Pro-resolving differentiation of macrophages and adaptive remodeling of the AVF

Postoperative inflammation initiates adaptive vascular remodeling but, when excessive or persistent, it may promote fibrosis and AVF failure.^16, 32^ We analyzed the temporal phenotypes of 15,901 mono/macs in the porcine arteries and veins searching for signatures that explained the high rates of physiological maturation in this model (**Figure 6A-B**). The postoperative infiltration of myeloid cells in both limbs of the fistula was confirmed by CD11b immunostaining (**Figure 6C**). Maximum myeloid infiltration was seen at 2 days postop. Interestingly, at baseline and 21 days, veins had significantly higher densities of CD11b+ cells than arteries from the corresponding time points. In both types of vessels, myeloid cells accumulated in the adventitia, supporting the role of the vasa vasorum in immune cell infiltration.

**Figure 6.**
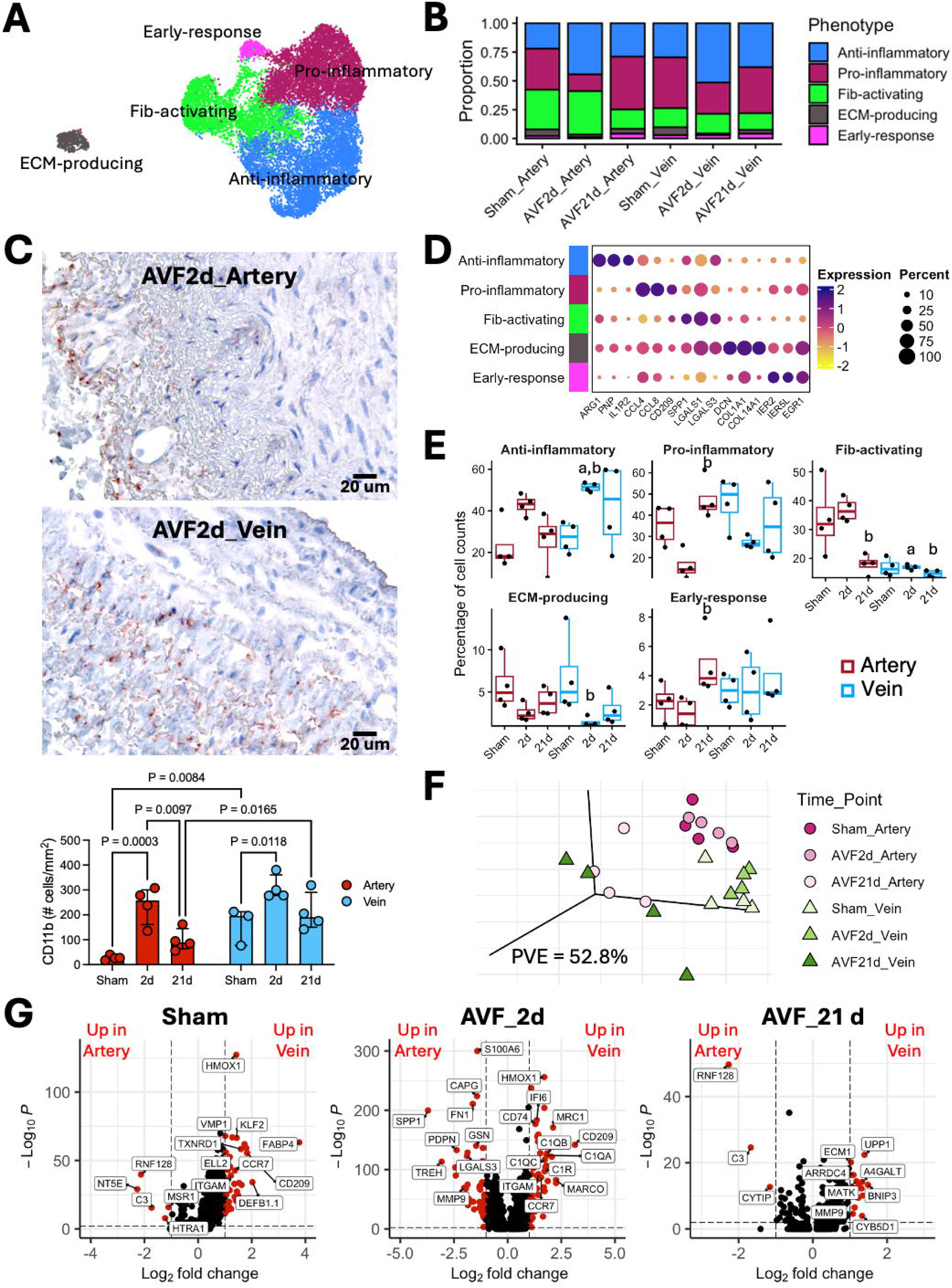
Regulation of postoperative inflammation in the pig AVF model. **A)** Integrated UMAP of monocyte/macrophages (mono/mac) from 12 femoral arteries and 12 femoral veins, color-coded by cell phenotype. **B)** Mean cell proportions of mono/mac phenotypes relative to the total number of mono/macs by experimental group. **C)** Representative immunohistochemistry and quantification of CD11+ cells in experimental groups. **D)** Dot plot representation of expression markers for mono/mac subpopulations. Dark purple dots indicate the highest expression levels, while the size of the dot represents the percentage of cells within each subcluster expressing the gene. **E)** Distribution of phenotypes per sample relative to the total number of mono/macs. a, P<0.05 compared with the artery; b, P<0.05 compared with the previous time point within the same vessel type. **F)** Principal component analysis of transcriptional variance in mono/macs at the sample level. **G)** Volcano plots of differentially expressed genes (DEG; FDR<0.01 and absolute log_2_(fold change [FC])>1) in mono/macs between arteries and veins by time point of tissue collection. Red dots indicate DEGs upregulated in arteries (log2FC<-1) or veins (log2FC>1).

Among mono/macs, we uncovered an anti-inflammatory subtype, defined by expression of arginase 1 (*ARG1*) and the decoy receptor *IL1R2*; pro-inflammatory mono/macs showing upregulation of chemokines and *CD209*; a fibroblast-activating phenotype over-expressing galectins (*LGALS1, LGALS3*) and osteopontin (*SPP1*); as well as minor subpopulations of early responders and ECM-producing macrophages (**Figure 6D**). To our surprise, and in contrast to the inflammatory mono/mac landscape observed in 7-day human AVFs,^16^ AVF creation in pigs induced an influx of anti-inflammatory mono/macs into both the arterial and venous walls while suppressing the expansion of pro-inflammatory subsets (**Figure 6E**). At 2 days, arteries also had higher abundance of fibroblast-activating macrophages than veins, but their proportion declined with remodeling, reaching levels comparable to those of veins by day 21. Venous macrophages also adapted to the change to arterial blood (**Figure 6F**). Heme oxygenase 1 (*HMOX1*), which is typically upregulated under hypoxic conditions,^33^ had higher expression in sham and 2-day veins compared with arteries, but not at 21 days (**Figure 6G**).

## DISCUSSION

Successful AVF maturation requires the coordinated enlargement of both the artery and the vein to provide sufficient blood flow for hemodialysis. Whether these two vascular beds adapt through shared or distinct molecular mechanisms has remained understudied. Using the first single-cell transcriptomic atlas of the porcine AVF, our analyses reveal that the postoperative adaptations of the artery and the vein occur in the setting of luminal endothelial dysfunction, expansion of the vasa vasorum, activation of pro-fibrotic fibroblast/myofibroblast programs, and controlled accumulation of macrophages. Despite their distinct developmental origins and specialized functions, both vascular beds progressively converge toward a shared transcriptional program during remodeling to promote the successful maturation of the AVF.

Convergent remodeling of the arterial and venous limbs is reflected in various cell populations of the AVF. These include the expansion of venous-like EC subpopulations likely in response to angiogenic stimuli, the temporal enrichment of anti-inflammatory macrophages, and the activation of common mechanosensitive and tissue repair programs in fibroblasts and myofibroblasts. Notably, this convergence was more pronounced in the artery than in the vein. By 21 days after AVF creation, the transcriptional landscapes of arterial ECs, pericytes, fibroblasts, and myofibroblasts more closely resembled their venous counterparts than their own preoperative states, supporting the concept of transcriptional arterial venification as a counterpart to the well-recognized process of vein arterialization. These findings argue against viewing AVF remodeling solely through the lens of vein maturation and instead support a model in which arteries and veins undergo parallel, yet incomplete, molecular reprogramming. It is likely that the molecular reprogramming of the artery after AVF creation primarily reflects an adaptive wound-healing response rather than a fundamental reconfiguration of its biomechanical properties. The latter are largely determined by the pre-existing ECM architecture and vessel wall composition, which may require substantially longer periods to undergo structural transformation. This delayed remodeling may help explain why inflow stenosis is considerably less common than outflow stenosis in newly created AVFs.

Another intriguing finding of this study was that the arterial and venous limbs underwent successful adaptive remodeling despite persistent endothelial denudation starting 2 days after anastomosis. These findings suggest that most shear stress sensing required to initiate vascular adaptation happens immediately after AVF creation. Similarly, human venous samples from both mature and failed second-stage AVFs collected at the time of transposition exhibit significant de-endothelialization irrespective of maturation outcomes.^16, 34^ Interestingly, both arteries and veins exhibited marked expansion of the vasa vasorum during remodeling, resulting in a similar diversity of microvessels in both types of vessels. This neovascularization process may provide an alternative route for NO delivery, nutrient exchange, and immune cell trafficking while the luminal endothelium remains compromised. Nevertheless, microvascular ECs in the venous wall retained inflammatory transcriptional programs that were not recapitulated in the artery, supporting the existence of a vessel-specific immune microenvironment, as previously described in human veins.^17, 18^ Together, these findings call for a reassessment of the long-held paradigm that successful AVF maturation depends on preservation of the luminal endothelium. Whether therapeutic strategies aimed at enhancing postoperative re-endothelialization or increasing NO bioavailability can further improve AVF maturation remains unknown.

Our findings also underscore the importance of circumferential wall stretching resulting from increased intraluminal pressure as a critical hemodynamic stimulus driving AVF remodeling. Although studies of biomechanical stretching have traditionally focused on SMCs,^35^ recent single-cell analyses of human AVFs have highlighted the central role of fibroblasts in vascular remodeling.^16^ Consistent with these observations, our temporal analysis revealed robust activation of mechanosensitive pathways in myofibroblasts and fibroblasts as early as 2 days after AVF creation in both the arterial and venous limbs. While additional cell populations may respond at earlier time points, the coordinated activation of these cell types strongly supports their role as orchestrators of ECM remodeling, cell migration, and tissue repair.

Finally, the porcine AVF exhibited several mechanisms that appear to promote successful maturation while limiting excessive postoperative fibrosis. These included the early predominance of anti-inflammatory macrophages, the progressive resolution of SPP1+ fibroblast-activating macrophages, and the coordinated induction of both profibrotic and antifibrotic regulators that may preserve ECM homeostasis. In contrast, human AVFs that fail to mature are characterized by the persistent presence of SPP1+ macrophages and sustained inflammatory macrophage-fibroblast crosstalk, which promote fibrotic remodeling.^16^

The limitations of the study include the lack of chronic kidney disease and the absence of maturation failure observed in patients. Despite these limitations, we provide the first high-resolution single-cell atlas of arterial and venous remodeling in a widely used pre-clinical animal model.^36^ This comprehensive molecular characterization enhances our understanding of AVF remodeling and the translational value of the model to accelerate the development of anti-stenotic therapies.

