## Supplementary Material for "The Venification of the Artery: Insights from Single-Cell Transcriptomics of a Porcine Arteriovenous Fistula Model"

### SUPPLEMENTARY MATERIALS AND METHODS

#### *Study Design*

Six Yorkshire pigs, ~50 kg in weight, were allocated to 1) bilateral femoro-femoral AVF creation for tissue harvest at 2 days postop, 2) bilateral AVFs for harvest at 21 days, or 3) bilateral sham operation to collect native vessels 2 days after opening and closing the skin and fascia. Each experimental arm included one male and one female. Therefore, a total of four arteries and four veins were collected per group for histology and single-cell RNA sequencing (scRNA-seq). The harvested AVFs included ~3 cm of each the artery and the vein from the juxta-anastomotic segment. The arterial and venous ends were separated at ~3 mm from the anastomosis; ~5-mm cross-sections distal to the anastomosis were stored in 10% neutral formalin for paraffin embedding, while the rest were enzymatically digested as described[1] to generate single cell suspensions.

#### *Surgical Procedures*

For creation of AVFs, pigs were pre-medicated with atropine (0.04 mg/kg), sedated with xylazine (2.2 mg/kg) and Telazol (4.4 mg/kg), and anesthetized using isoflurane. All surgical procedures were performed by board-certified vascular surgeons (M.S.S., M.T.). Bilateral femoral cutdowns were performed just below the inguinal ligament, and fascial and muscle planes were carefully dissected out with attention to hemostasis. The femoral artery and vein were identified and skeletonized. The vein was transected using silk ligature and the proximal portion was occluded with a bulldog clamp. The artery was controlled with bulldog clamps. The AVF was created using an ~5 mm end-to-side anastomosis with 7-0 polypropylene suture. Vessel clamps and loops were then released, with special attention to obtaining hemostasis and ensuring there was no kinking or torsion of the AVF. This was done in identical fashion bilaterally. The fascia and skin were then closed in layers using absorbable sutures. Intramuscular buprenorphine (0.03 mg/kg) was administered postoperatively followed by a fentanyl patch on the back dosed at 25-50 ug/hour for 3 days. Blood flows were measured at the time of AVF harvest using a portable Philips Lumify ultrasound (Philips, Amsterdam, Netherlands). Animal experiments were approved by the University of Miami Institutional Animal Care and Use Committee.

*Single-Cell RNA Sequencing*

Sequencing was performed at the University of Miami John P. Hussman Institute for Human Genomics. Briefly, samples with >80% viability were run using the Chromium Single Cell 3′ Library & Gel Bead Kit v3 (10X Genomics, Pleasanton, CA). Each sample was processed on an independent Chromium Single Cell A Chip (10X Genomics) with a target capture of 10,000 cells. Sequencing libraries were evaluated for quality on the Agilent Tape Station (Agilent Technologies, Palo Alto, CA), quantified using a Qubit 2.0 Fluorometer (Invitrogen, Carlsbad, CA), and qPCR before sequencing on the Illumina NovaSeq 6000. FASTQ files were generated with Cell Ranger's mkfastq pipeline (version 6.0.2). The Cell Ranger's count pipeline (version 6.0.2) was used to generate raw gene-barcode matrices from alignment to the *Sus scrofa* 11.1 reference genome. Sequences were analyzed in Seurat v5 and integrated with Harmony using published bioinformatic pipelines.[2-4] Compositional analyses were performed using the Wilcoxon test in SeuratExtend.[5] The swine sequencing data were deposited in accession number GSEXXXXXX.

*Histology and Immunostaining*

Tissue sections were stained with hematoxylin and eosin for gross morphometric analysis. Average wall thickness was quantified as the mean of four equidistant measurements. Intimal hyperplasia was quantified as average intimal thickness and intima/media (I/M) area ratio using ImageJ (National Institutes of Health, Bethesda, MD). Cell density was calculated as the number of cells divided by wall area using 4′,6-diamidino-2-phenylindole (DAPI) stained sections. Cell counting was performed in Image-Pro v11.1 (Media Cybernetics, Rockville, MD). Specific cell markers were stained by immunohistochemistry (IHC) or immunofluorescence (IF). Briefly, paraffin sections were rehydrated by serially immersing them in xylene, alcohol, and water, followed by antigen retrieval in 10 mM sodium citrate buffer, pH 6.0 or Tris-EDTA buffer, pH 9.0 in a pressure cooker (106-110$^{o}$C) for 10 min. For IHC, sections were treated with Dako Peroxidase Blocking Reagent (Dako-Agilent, Santa Clara, CA) for 10 min, Dako Protein Block for 1 hour, and primary antibody (Ki-67, 1:25, pH 6, #550609; BD Pharmingen, San Jose, CA; CD11b, 1:100, pH 9, #ab133357; Abcam, Waltham, MA) overnight at 4$^{o}$C in Dako Antibody Diluent. Bound antibodies were detected by incubating with anti-mouse Dako EnVision+ Single Reagent (#K4001) for 1 hour, followed by AEC substrate (#ab64252, Abcam) for 10 min. Sections were counter-stained with hematoxylin and mounted in aqueous mounting media.

For IF, after antigen retrieval, slides were incubated with TNB Blocking Buffer (#FP1012; Akoya Biosciences, Marlborough, MA) for 1 hour, followed by primary antibodies diluted in TNB overnight at 4$^{o}$C (CD31, 1:25, pH 6, #ab28364, Abcam; eNOS, 1:100, pH 6, #LS-C413412-30; LS Bio, Seattle, WA). Bound antibodies were detected with Alexa Fluor 546 goat anti-rabbit antibody (1:1000, #A11081; Thermo Fisher Scientific, Waltham, MA) for 45 minutes. Sections were counter-stained with 300 nM DAPI solution (#D1306, Thermo Fisher Scientific) in PBS for 5 minutes and mounted in fluorescence-compatible mounting medium (#ab104135, Abcam). Sections were examined in a Keyence All-in-One Fluorescence Microscope BZ-X800L and photographed using the Keyence BZ-X800 Viewer software (Keyence, Itasca, IL).

*Statistical Analyses*

Statistical analyses were performed using GraphPad Prism 10.1.1 (San Diego, CA). Normally distributed data (Shapiro-Wilk test) were compared using t-tests and expressed as mean ± standard deviation. A two-way ANOVA was performed for multigroup comparisons, with uncorrected Fisher's LSD post hoc test and matching of arteries and veins from the same AVF or anatomical site for shams. If normality assumptions were not met, the Mann-Whitney test was used, and data were expressed as median and interquartile range. A Friedman test was used for nonparametric multi-group studies, with uncorrected Dunn post hoc test. Statistical significance was defined as $P<0.05$.

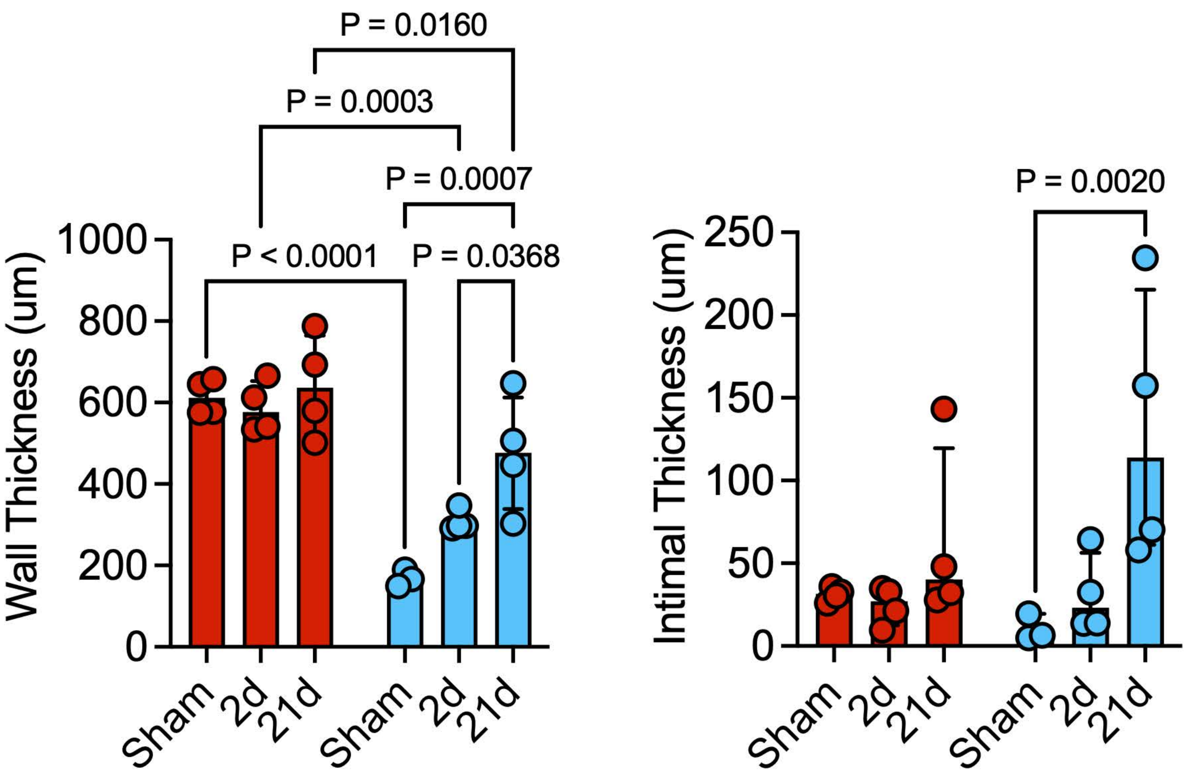


**Supplementary Figure S1.** Temporal histomorphometry of arteries and veins in the pig AVF model. Error bars indicate the median and interquartile range. Only significant p-values are shown.

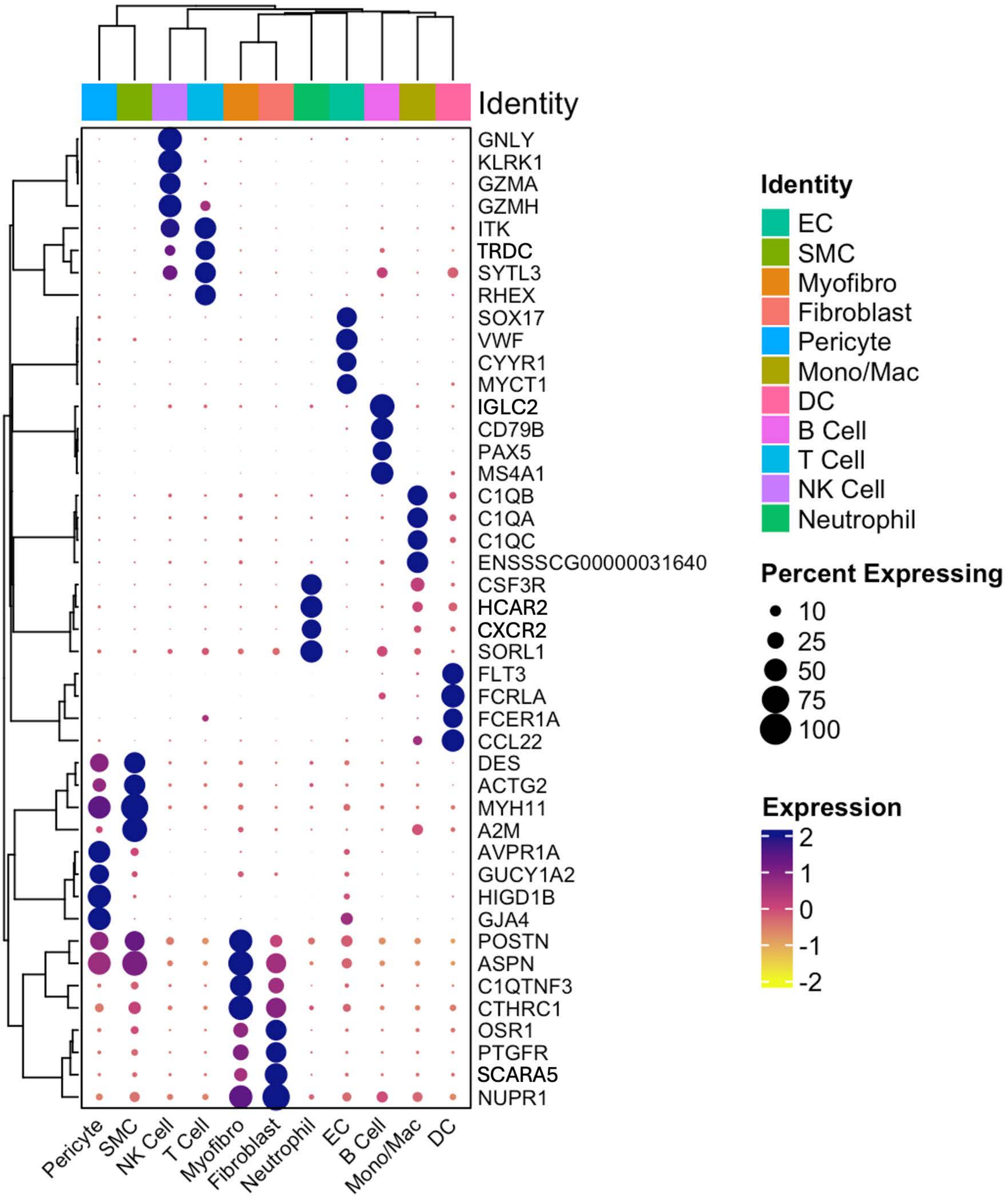


**Supplementary Figure S2.** Expression markers for the main cell subpopulations in the integrated single-cell map. Dark purple dots indicate the highest expression levels, while the size of the dot represents the percentage of cells within each cluster expressing the gene.

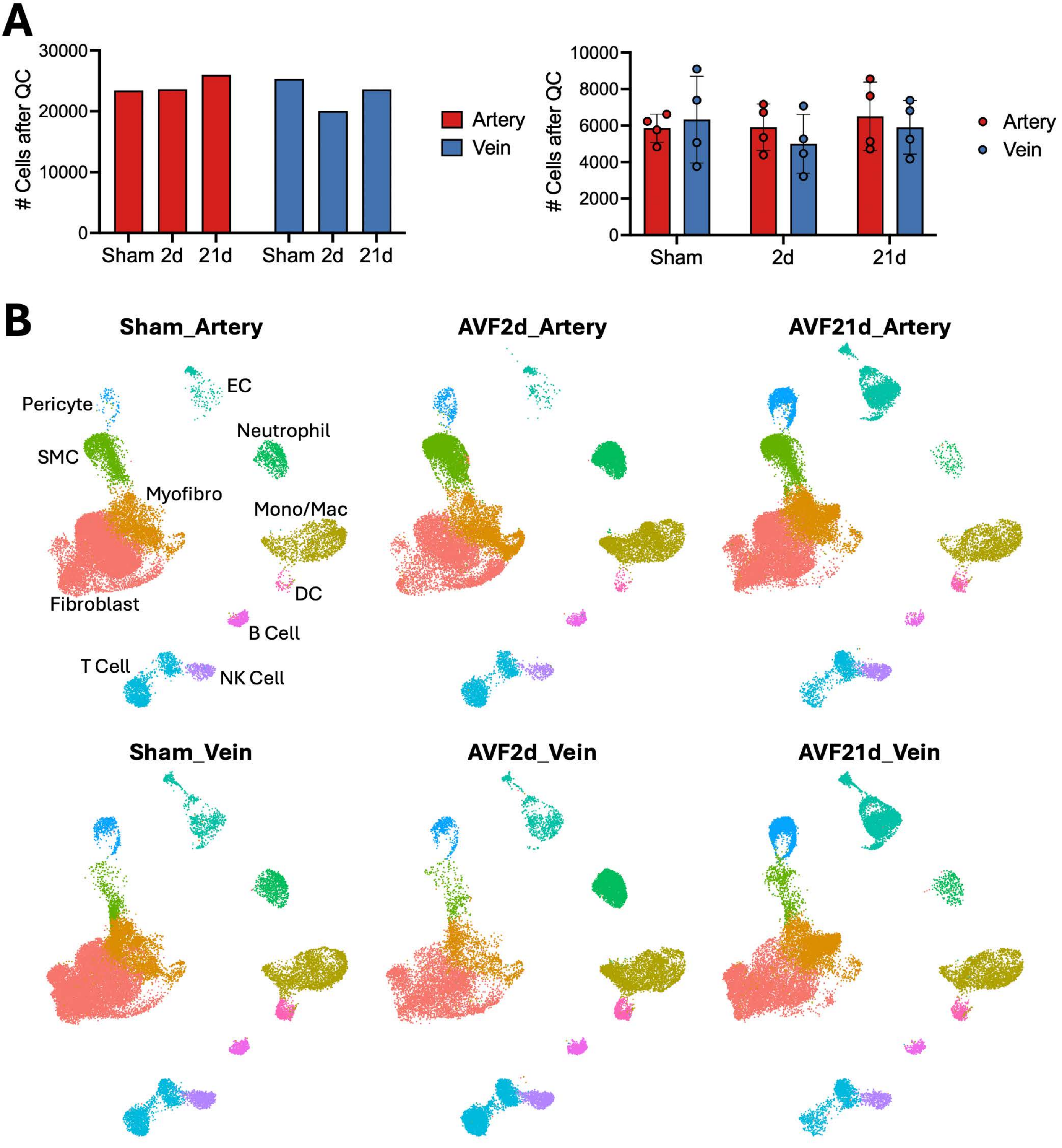


**Supplementary Figure S3. A)** Quality control (QC) of single-cell libraries per experimental group and individual sample. **B)** Uniform manifold approximation and projections (UMAPs) of cells isolated per experimental group.

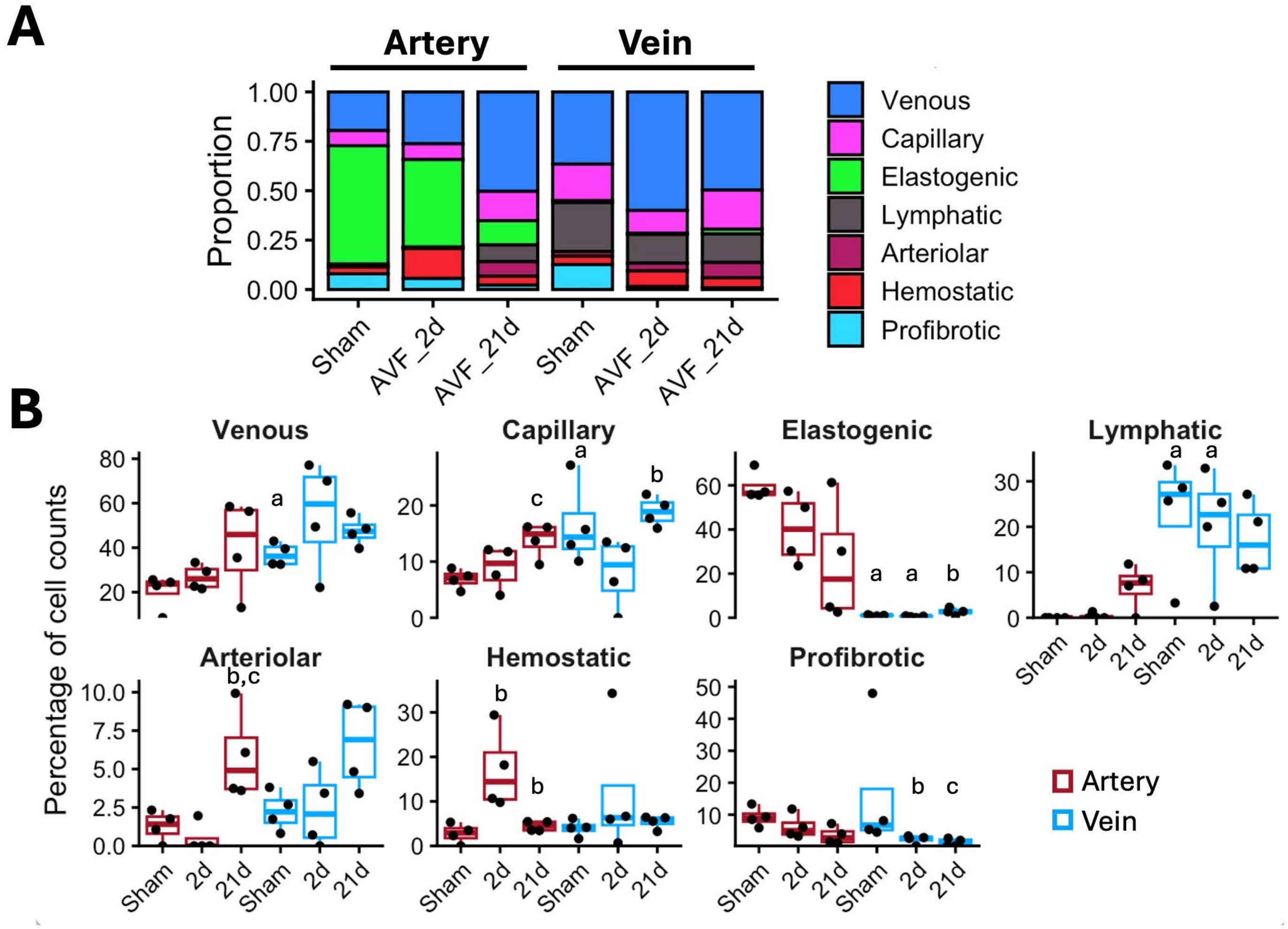


**Supplementary Figure S4. A)** Mean cell proportions of endothelial cell (EC) phenotypes relative to the total number of ECs by experimental group. **B)** Relative proportions per sample. a, $P<0.05$ compared with the artery; b, $P<0.05$ compared with the previous time point within the same vessel type; c, $P<0.05$ compared with the sham group within the same vessel type.

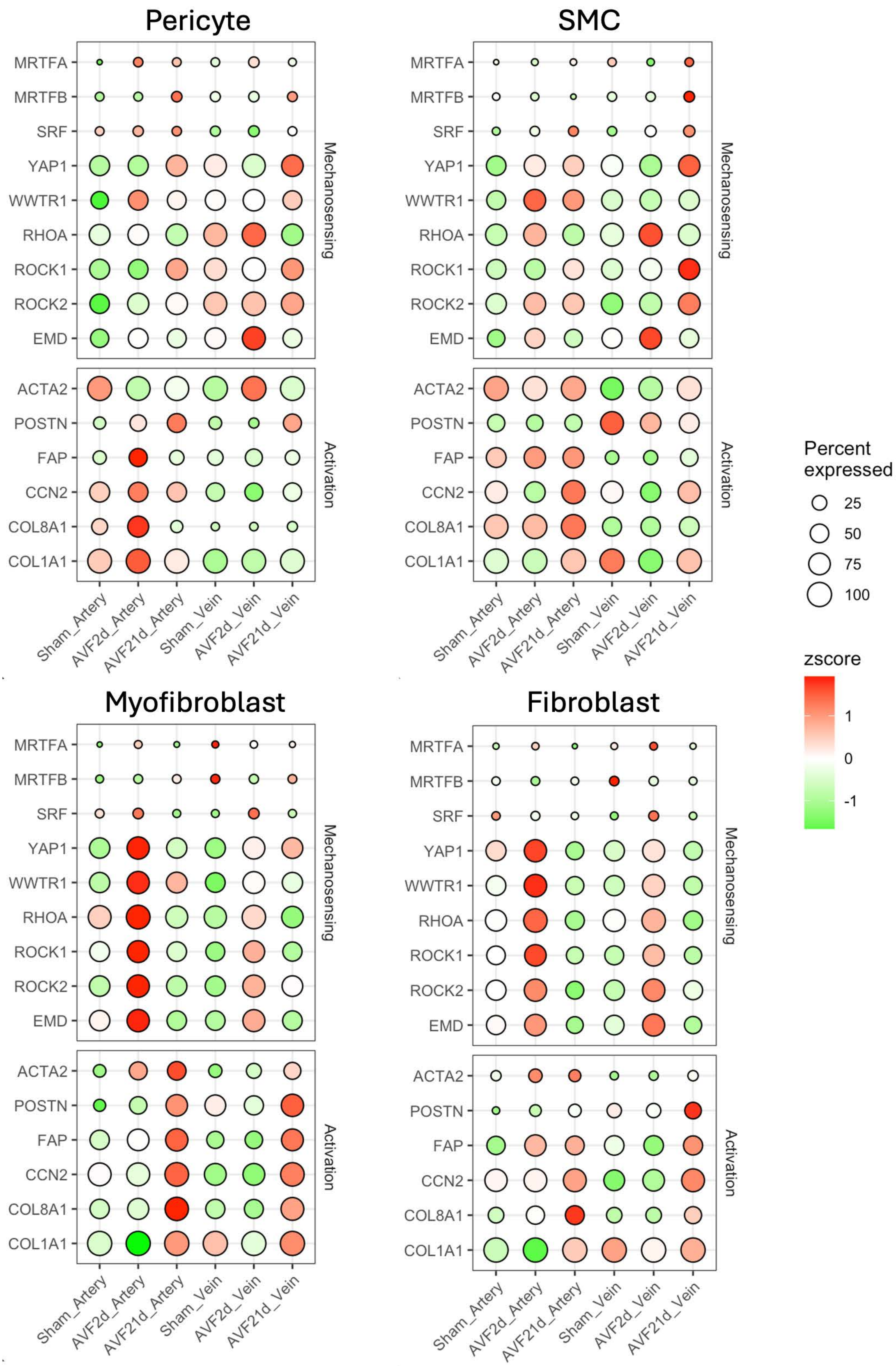


**Supplementary Figure S5.** Time-dependent upregulation of mechanotransduction genes and markers of fibroblastic cell activation in arteries and veins. Bright red dots indicate the highest expression levels, while the size of the dot represents the percentage of cells expressing the gene.

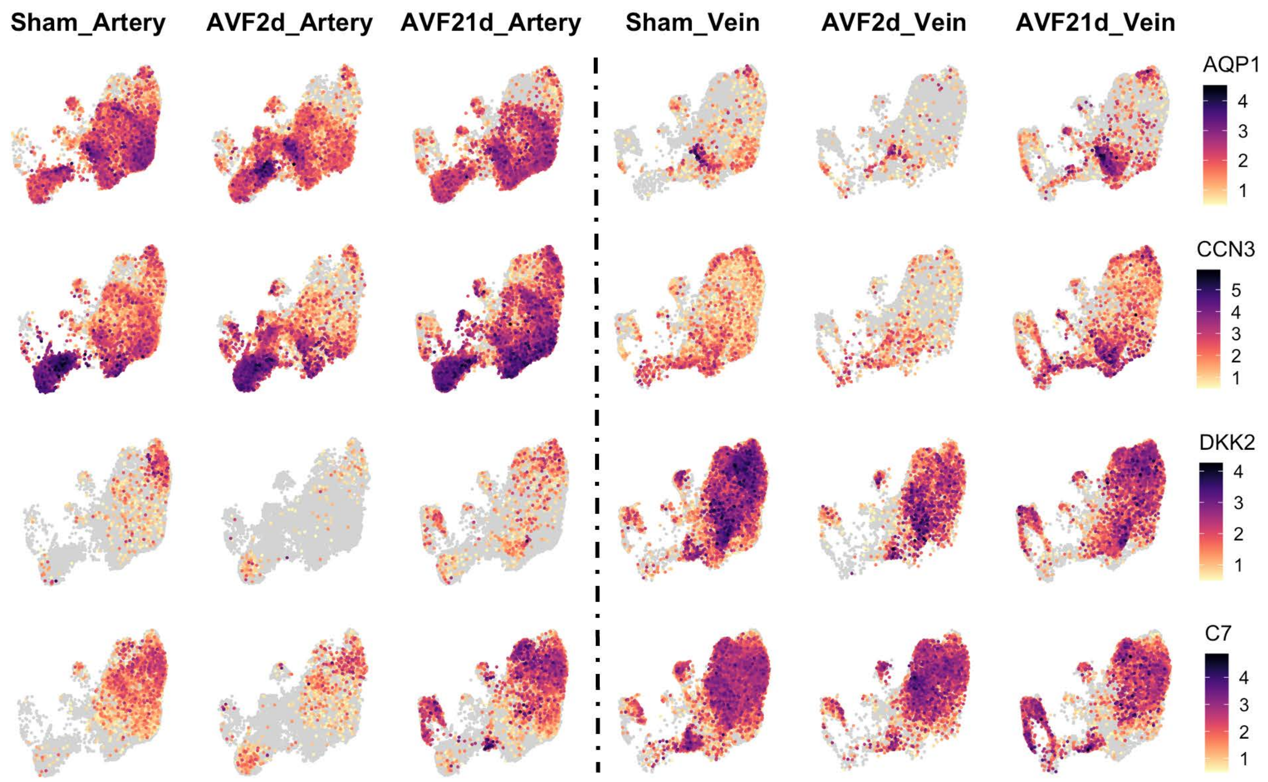


**Supplementary Figure S6.** Differentially expressed genes between arteries and veins, independent of the time point, suggest inherent vessel-selective mechanisms of myofibroblast and fibroblast differentiation. Focused UMAPs of mural cells, myofibroblasts, and fibroblasts from the different experimental groups with cells color-coded according to the expression levels of the genes to the right. The gray background of the maps indicates no expression detected.
